# Pseudo-time trajectory of single-cell lipidomics: Suggestion for experimental setup and computational analysis

**DOI:** 10.1101/2025.04.11.648323

**Authors:** Paul Jonas Jost, Daniel Weindl, Klaus Wunderling, Christoph Thiele, Jan Hasenauer

**Affiliations:** Bonn Center for Mathematical Life Sciences, University of Bonn; LIMES Life and Medical Sciences Institute, University of Bonn, Bonn, Germany

## Abstract

Cellular heterogeneity is a fundamental facet of cell biology, influencing cellular signaling, metabolism, and gene regulation. Its accurate quantification requires measurements at the single-cell level. Most high-throughput single-cell technologies provide only a snapshot of cellular heterogeneity at a specific time point because the measurement is destructive. This limits our current ability to understand the dynamics of cellular behavior and quantify cell-specific parameters.

We propose an experimental setup combined with a model-based analysis framework, enabling the extraction of longitudinal data from a single destructive measurement. Although broadly applicable, we focus on lipid metabolism, a domain where obtaining longitudinal single-cell data has remained elusive due to technical constraints.

Our method leverages multiple labels whose measurements are linked to a shared dynamic. This allows the estimation of cell-specific parameters and the quantification of heterogeneity. This framework establishes a foundation for future investigations, providing a roadmap toward a deeper understanding of dynamic cellular processes.

## Introduction

Cell heterogeneity, the inherent variability among individual cells within a seemingly homogeneous population (Figure 1A), has emerged as a focal point in recent scientific research [Lee et al., 2020, Muto et al., 2021, Kinker et al., 2020]. It can be analyzed based on measurements of a large number of biochemical species through techniques such as single-cell transcriptomics [Tang et al., 2009], single-cell proteomics [Kelly, 2020], and single-cell lipidomics [Thiele et al., 2019]. Complementary to this, single-cell imaging technologies have been developed for the longitudinal analysis of a few selected biochemical species [Skylaki et al., 2016]. These technologies had a profound impact across various domains, including cancer research [Hosein et al., 2019, Yabo et al., 2021], immunology [Schnell et al., 2023], microbiology [Hatzenpichler et al., 2020], and adipocyte physiology [Wang et al., 2021].

**Figure 1:**
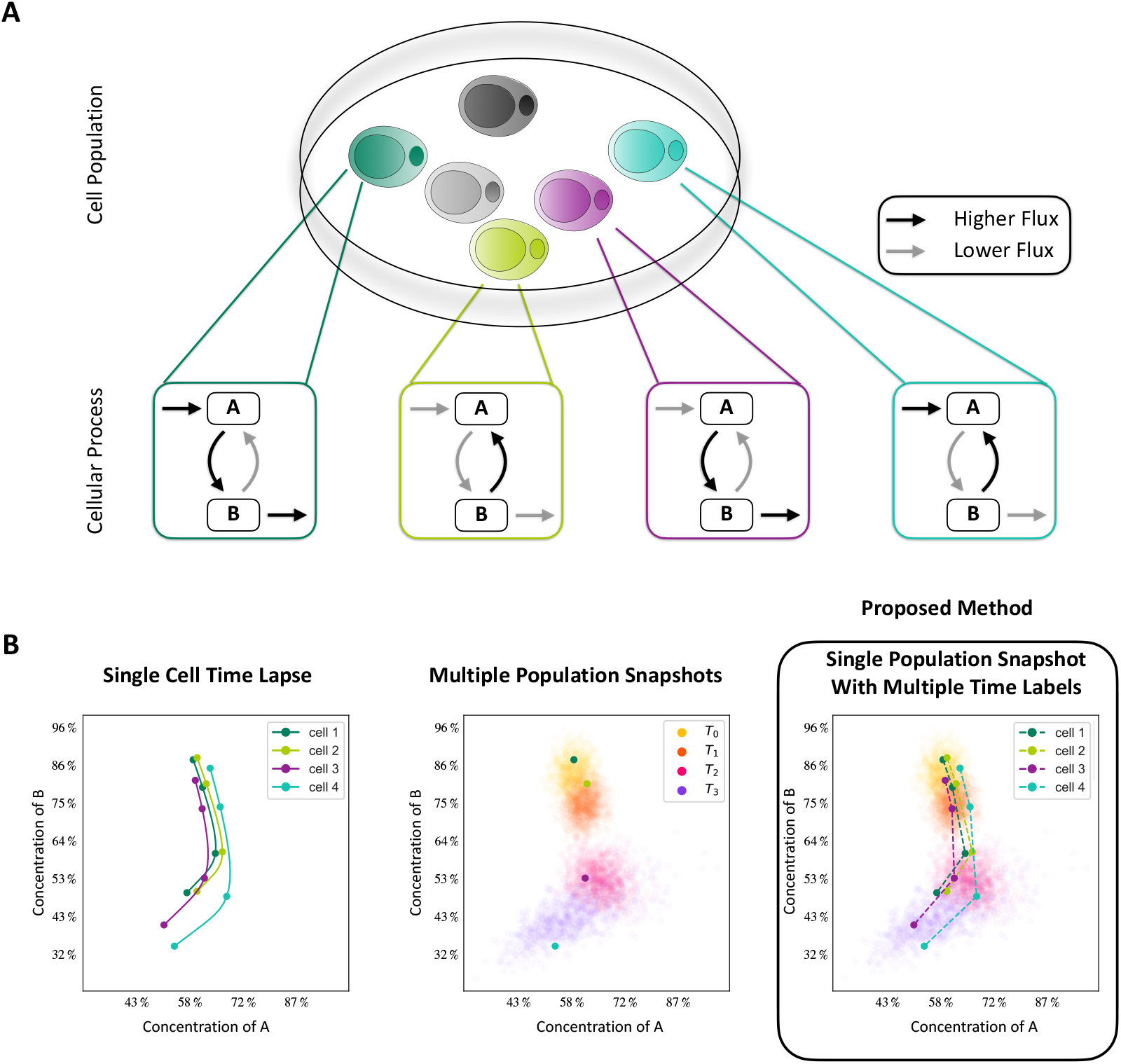
Cell heterogeneity and different types of measurements. (A) Different cells may differ in their cellular process activities, preventing two distinct cells from being used as measurements at two different time points. (B) To capture the time-lapse of single cells (left), single-cell destructive measurements at different time points can only yield information on the distribution over time and no accurate dynamics (center). Our proposed method resolves this issue, allowing for accurate dynamics of a single cell as well as distributions for the whole population (right).

Most single-cell studies use inherently destructive measurement techniques, which provide information about the state of cells at the time of the measurement [Kelly, 2020, Thiele et al., 2019, Hosein et al., 2019]. However, dynamic processes within cells and differences in signal processing, gene regulation, and metabolism mostly remain elusive (Figure 1B). This is problematic as most cellular processes are inherently dynamic, and small changes in rates of reactions can result in large differences, including different fate decisions.

In the field of single-cell transcriptomics, dynamic processes are starting to be addressed by distinguishing mRNAs at different processing stages [Bergen et al., 2021] or metabolic labeling of mRNAs [Erhard et al., 2022]. These approaches provide information about the time point of transcription, enabling velocity calculations and pseudo-time analysis [Erhard et al., 2022]. Although metabolic labeling is well established in bulk proteomics [Lindemann et al., 2017, Alabert et al., 2020] and lipidomics [Kuerschner and Thiele, 2022, Triebl and Wenk, 2018], on the single-cell level, they currently have no velocity or pseudo-time analysis methods available.

The spectrum of available metabolic labeling techniques is broad. Two important labeling strategies for lipids are stable isotope labeling [Allen et al., 2015] and alkyne-fatty acid click labeling [Kuerschner and Thiele, 2022]. Click-labeling allows reporter molecules to be linked to their targets so that the different labeled species can be easily distinguished, for example, by fluorescence microscopy or mass spectrometry. Different reporters can be clicked on different samples, which are then combined into one measurement to reduce experimental variability. The technique has been successfully utilized with mass spectrometry-based lipidomics to assess population averages [Thiele et al., 2019, Wunderling et al., 2023]. Moreover, the study of single cells has been demonstrated [Thiele et al., 2019], but the destructive nature of the measurement hinders the assessment of single-cell trajectories.

This study proposes an experimental protocol paired with a model-based analysis framework to assess dynamic lipid processing within individual cells based on a single destructive measurement. The protocol includes using multiple metabolic labels introduced at different time points prior to the measurement. Under the assumption that the metabolic labeling does not influence metabolism, we attribute the measurements for the different labels to a joint dynamic and thus provide what can be interpreted as multiple measurements along a trajectory. The information in the measurements is then deconvoluted using a knowledge-based dynamic model of lipid metabolism. We show that using this setup, a single destructive measurement can provide information about the state of the dynamic lipid processing in individual cells. Overall, the approach shares similarities with isotopically nonstationary metabolic flux analysis [Cheah and Young, 2018] but allows for the analysis of cell-to-cell variability. We use synthetic data as a proof of concept and show that we can accurately recover single-cell parameters, allowing us to quantify cell heterogeneity.

## Results

### Model-based approach for assessing cellular dynamics based on a single measurement

To assess the cell-to-cell variability of metabolic processing, we propose a simple experimental protocol paired with a model-based analysis framework. The core idea is to provide cells with differently labeled metabolites at different time points and to measure the concentration of the metabolites at a single time point. As the labeled metabolites have remained in the system for varying amounts of time, the labels implicitly provide a time stamp. The model-based approach enables the assessment of the dynamical properties of individual cells by leveraging this implicit information.

We focus on a specific setup for illustrative purposes. We consider cells in a growth medium, which at specific time points are susceptible to changes within this medium. Specifically, the only perturbation within the medium is the introduction of different labels at different time points. We make two additional assumptions: (1) unlabeled and labeled metabolites possess the same biochemical properties – meaning that reactions taking place within cells are agnostic to labeling –, and (2) the metabolite levels (disregarding the label) in the medium are considered to be constant throughout the experiment. In such a setup, the label itself, as well as the availability of labeled metabolites, does not perturb the system, and it will remain in a metabolic steady state, meaning that for any metabolite, the sum over all its unlabeled and labeled forms will stay constant.

The experimental setup and the dynamics of a metabolite are illustrated in Figure 2A. In the case of three labels, the setup is as follows:

**Figure 2:**
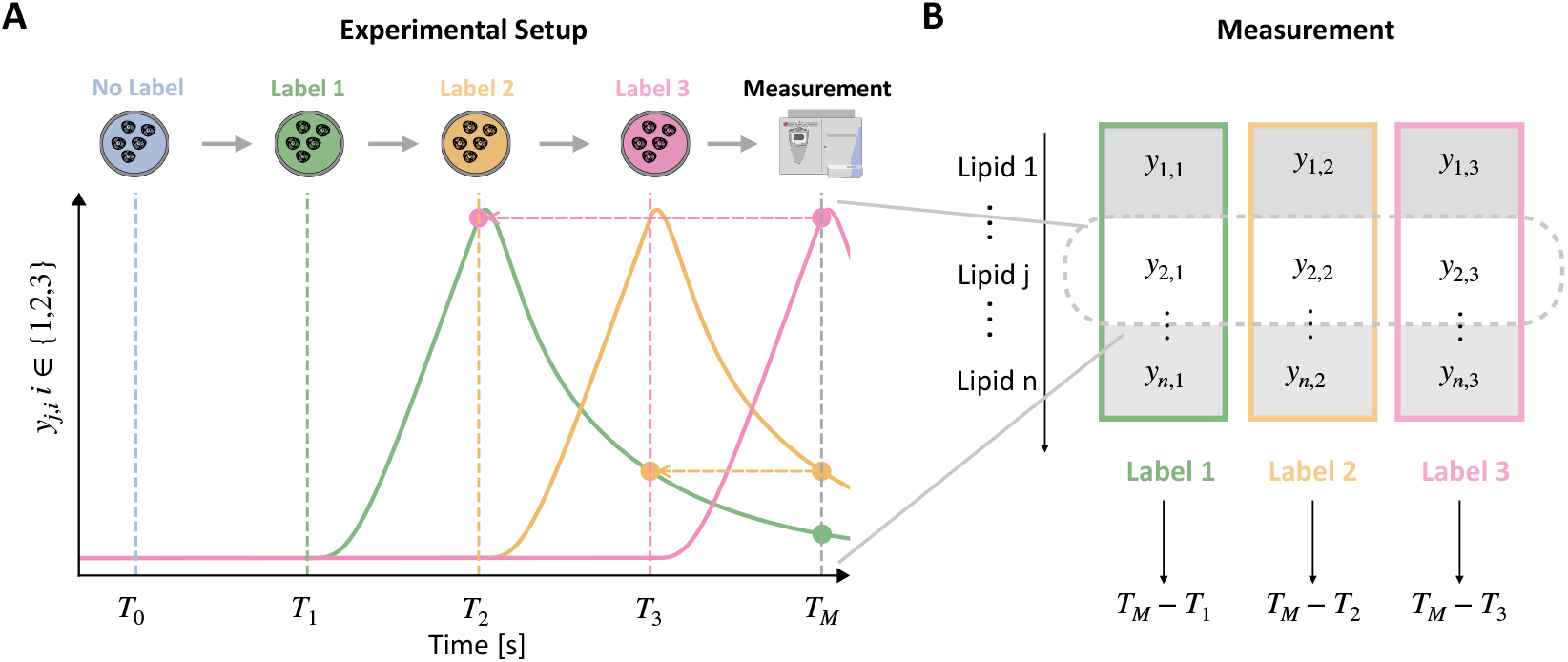
Illustration of proposed experimental setup and measurement. (A) Experimental setup paired with exemplary dynamics of a labeled chemical species for the case of three distinct labels. The experiment starts in the steady state for the label-free growth medium, and then growth media containing labeled species are provided after washing the cells (much like a pulse-chase experiment). The growth media only differ in the label provided and are otherwise identical. This setup results in three time-shifted trajectories of the differently labeled species. (B) The cells are harvested, and label-sensitive measurements provide a pseudo-time trajectory.

(Step 0) Cells are incubated with label-free growth medium at time point *T*_0_ for a sufficiently long period to allow the cells to reach a steady state.

(Step 1) At time point *T*_1_ (≫ *T*_0_), cells are washed, and a growth medium containing a metabolite with label 1 is applied.

(Step 2) At time point *T*_2_ (> *T*_1_), cells are washed, and a growth medium containing the same metabolite with label 2 is applied.

(Step 3) At time point *T*_3_ (> *T*_2_), cells are washed, and a growth medium containing the same metabolite with label 3 is applied.

(Step 4) At time point *T*_*M*_ (> *T*_3_), cells are harvested and concentrations of the differently labeled (and potentially also the label-free) metabolites are measured.

The experiment can be performed with any number of labels *N*_*L*_ ≥ 1. As we will discuss, while more labels provide more information about individual cells, they also increase the complexity of the experiment and subsequent analysis.

Notably, under the experimental conditions outlined above, the concentrations of the metabolites that possess only label *l* are time-shifted compared to the metabolites with label *l*^′^. The time shift is equal to the difference between the time points at which the respective labels were applied to the medium, i.e., 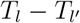 (Figure 2B). A mathematical proof for this core result is provided in the Supplementary Material. The proof of time-shifting confirms that one measurement at a single time point can provide information about a time course if a cell is exposed to a sequence of growth media with different labels. However, extracting the longitudinal information encoded in such measurements requires a label-aware dynamical model, which can easily be generated from a rule-based framework.

In the following, we will show how the proposed experimental setup can be used to assess cell-to-cell variability in metabolic processes and to determine causal differences between cells. Therefore, we assume that experimental data of the proposed setup are available for *n*_*c*_ cells of a heterogeneous cell population, yielding datasets 𝒟_*j*_ with *j* = 1, …, *n*_*c*_. We complement this with dynamic modeling to quantify the cell-to-cell variability. The mathematical model is a set of ordinary differential equations based on the biochemical reactions that are part of the considered metabolic process,

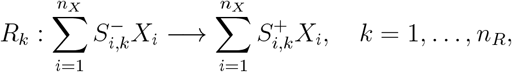

with biochemical species *X*_*i*_ and stoichiometric coefficients 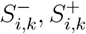.The reaction rates are modeled using mass-action kinetics. To account for the inherent cell-to-cell variability, we allow the vector of reaction rate parameters, 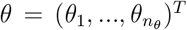, to differ between cells. A common assumption is that it is sampled from a probability distribution, *θ* ∼ *p*(*θ*) (similar to [Hasenauer et al., 2011] and related studies), e.g., a multivariate log-normal distribution [Fröhlich et al., 2018b].

Given the single-cell datasets and the mathematical model, assessing the (source of the) cell-to-cell variability is equivalent to determining the cell-specific parameters *θ*^(*j*)^ from 𝒟_*j*_, for *j* = 1, …, *n*_*c*_. In this study, we solve this problem using a maximum likelihood approach. Parameter optimization is performed using a multi-start local optimization scheme, while parameter uncertainty analysis is performed using the Fisher information matrix (see Methods section and Supplementary Material).

### Mathematical model for fatty acid metabolism in adipocytes

To assess the possibility of the reconstruction of single-cell dynamics and parameters from single-cell snapshot measurements, we performed a comprehensive *in silico* study. In our model, we consider fatty acid metabolism in adipocytes. This model is suitable from both a technical and biological standpoint. Technically, adipocytes have high lipid content, making it easier to pass the sensitivity threshold of mass spectrometers. Biologically, adipocytes play a key role in energy storage [Rosen and Spiegelman, 2014] and are very heterogeneous [Kahn et al., 2019, Luong et al., 2019].

For the *in silico* study, we constructed a model for the core components of fatty acid metabolism (Figure 3A). The model accounts for triacylglycerol (TAG) synthesis and lipolysis [Large et al., 2004], accordingly, including glycerol (GLY) and glycerol-3-phosphate (G3P), lysophosphatidic and phosphatidic acids (LPA, PA) as well as mono-, di-, and triacylglycerols (MAG, DAG, TAG). Additionally, we include the phospholipids phosphatidylcholine (PC), phosphatidylethanolamine (PE), and phosphatidylserine (PS), which are key components of cell membranes and in cell signaling. Additional metabolites of the phospholipid metabolism were also incorporated. We simplify the model by assuming constant ATP and CoA levels, subsequently omitting them from the model. Thus, we use fatty acids directly in reactions involving acyl-CoA. Overall, the model accounts for 22 metabolites and 40 reactions. A comprehensive list is provided in Supplementary Tables S1 and S2. The reaction rates are assumed to follow mass action kinetics. Hence, each reaction possesses one parameter. Those parameters are generally unknown, and we use dynamic modeling to estimate them. Cell-to-cell variability can be quantified through their differences across cells.

**Figure 3:**
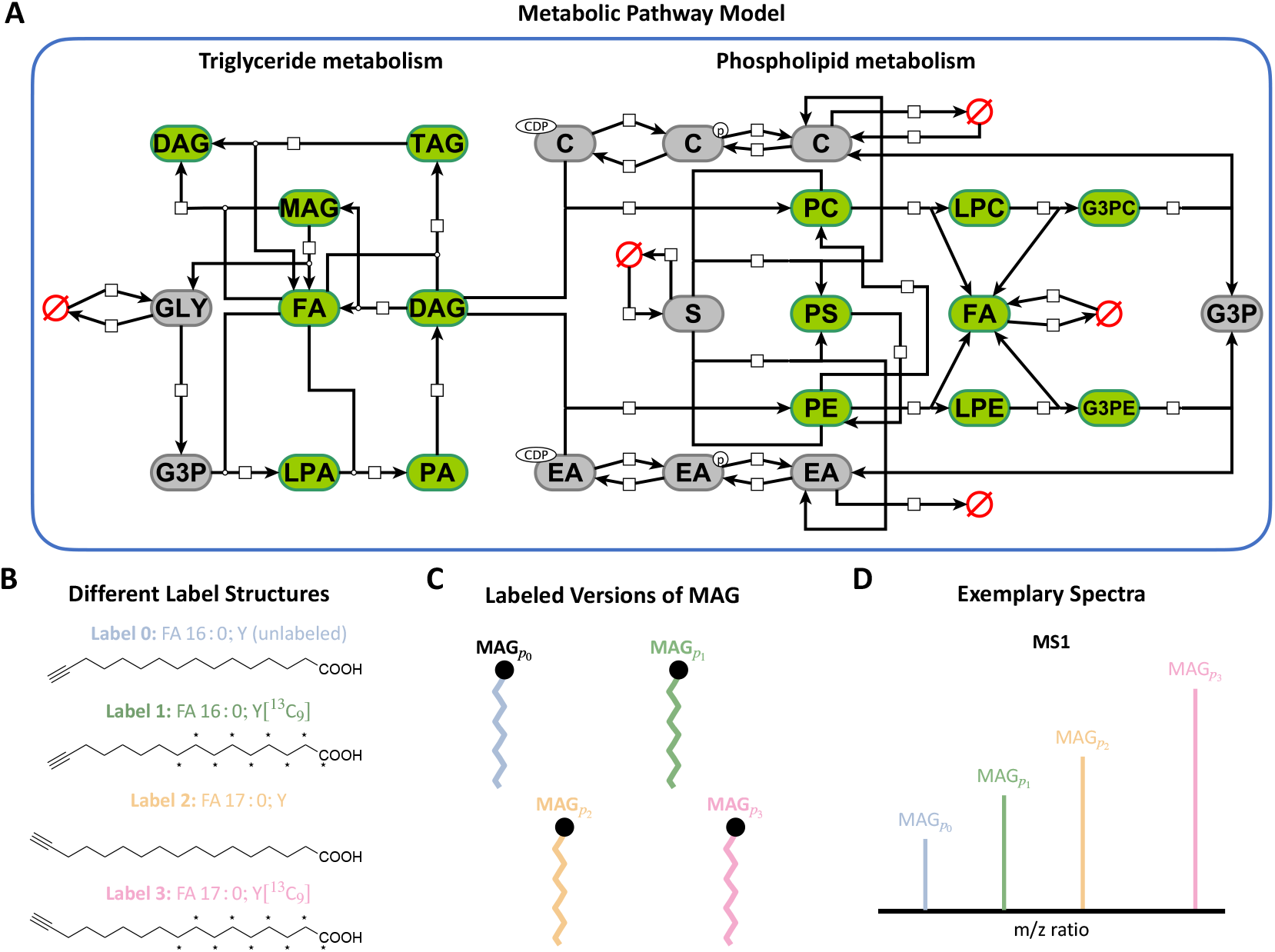
*In silico* mechanistic model of lipid metabolism and metabolic labeling setup. (A) Visualization of the considered model of lipid metabolism in adipocytes, considering the two main parts of Triglyceride and phospholipid metabolism (Visualization following SBGN [Novère et al., 2009] notation). Species with green boxes are observed, while species with grey boxes are not observed but are still considered in the model. Red empty sets symbolize the in- and outfluxes of the model. (B) Chemical structures of the three alkyne-fatty acids FA 16:0;Y, FA 16:0[^13^C_9_];Y, and FA 17:0;Y, which serve as labels in the experimental setup. (C) Illustration of the species MAG with different labels. A more extensive example can be found in Supplementary Figure S1. (D) Illustration of the mass spectrometry measurement. The labels are chosen so they can already be distinguished in the mass spectrometer.

As the lipid metabolism includes and processes extracellular fatty acids (FAs), we considered an *in silico* experimental setup in which cells are provided with alkyne-fatty acids (FAs) functioning as labels (Figure 3B). FAs are available in different chain lengths and with different biochemical properties, but several variants with similar chain lengths possess comparable biochemical properties [Raclot, 2003]. Thus, we use the alkyne-fatty acids FA 16:0;Y, FA 16:0[^13^C_9_];Y, and FA 17:0;Y as labels, which can be distinguished using mass spectrometry to determine the label-resolved concentrations of lipids or groups of lipids (Figure 3D). In line with this, the model accounts for multiple labels that do not alter the biochemical properties of the molecules but can be distinguished in the measurement process (Figure 3C, D). A biochemical species, such as DAG and TAG, might carry multiple fatty acids-hence, multiple labels are possible. For these species, the number of considered state variables corresponds to the number of possible label combinations. We assume that the position of the label can be disregarded, reducing the number of possible combinations (see Supplementary Information for details on the model formulation).

The resulting dynamical model for fatty acid metabolism in adipocytes translates for the case of three labels to an ordinary differential equation (ODE) model with 101 metabolites and 263 reactions. The number of metabolites and reactions grows quickly with the number of labels. In contrast, the number of parameters remains constant as the labels are assumed not to affect the processing of metabolites. Considering every metabolite that contains one or more fatty acids within the model as observable results in a total of 92 observables. For the subsequent assessment, we used this model to generate quantitative noise-corrupted data for different numbers of labels.

### Optimization allows for reliable fitting of dynamical models

To evaluate the feasibility of using a model-based approach to assess experimental data collected under the proposed protocol, we conducted an extensive *in silico* study (Figure 4A). Utilizing the mathematical model for fatty acid metabolism in adipocytes, synthetic data were generated for a heterogeneous population comprising *n*_*c*_ = 1000 cells (see Method section for parameter distribution details), providing a ground truth as a reference for evaluation.

**Figure 4:**
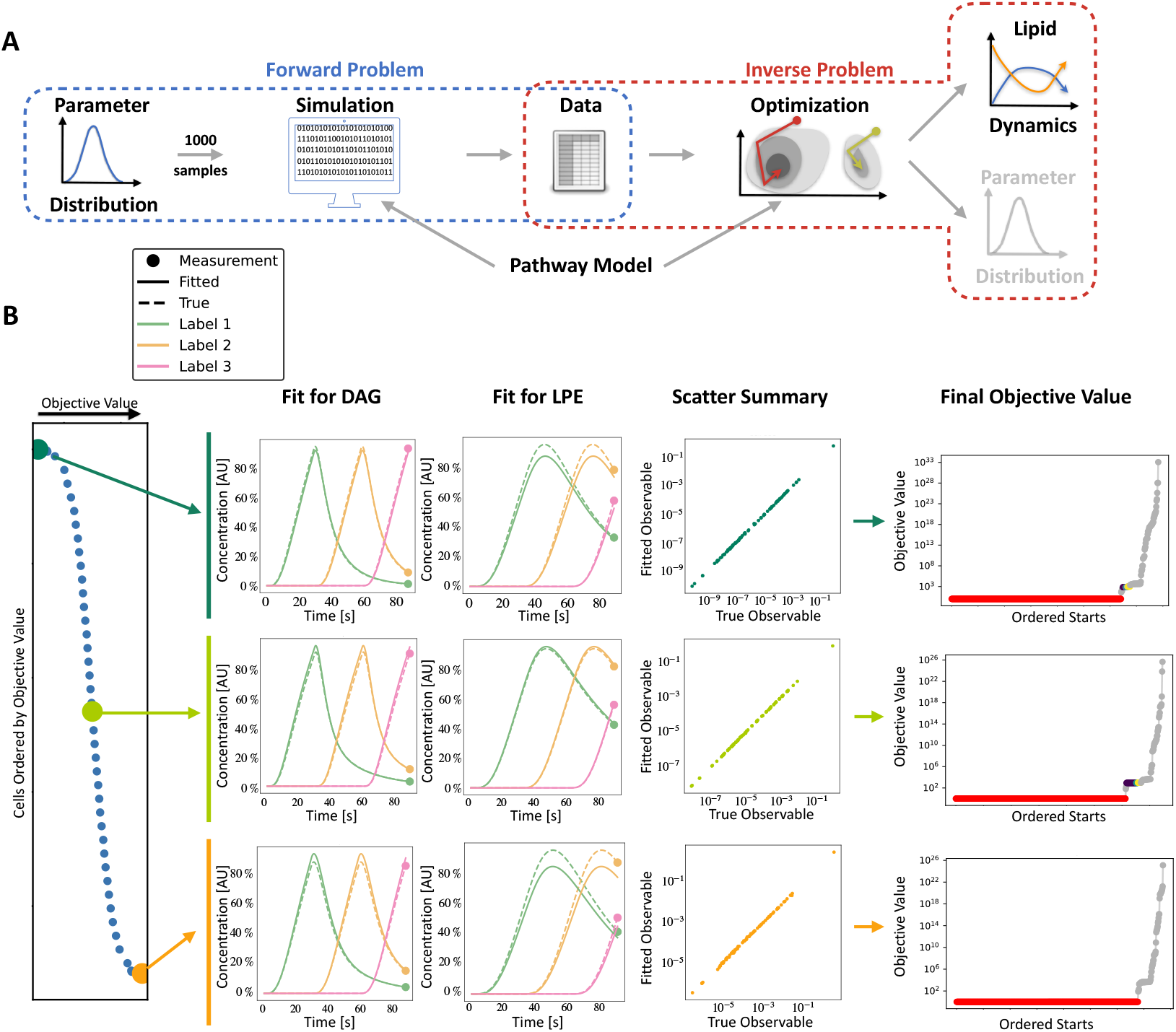
Evaluation setup and assessment of the parameter optimization pipeline. (A) Schematic overview of the workflow: In the forward problem, parameter vectors for individual cells are sampled independently from a multivariate log-normal distribution. For each parameter vector, the model is simulated. The resulting simulated values for each observable are corrupted by adding normally distributed noise. In the inverse problem, for each synthetic dataset, a multi-start local optimization is performed, and the results are analyzed in terms of goodness of fit. The pathway model is used in the forward and inverse problems. (B) Illustration of fitting results for synthetic datasets: (left) Ordering of cells by objective function value, with smaller values indicating a better fit. (Middle) Best fits for the two selected species, DAG and LPE, for best, median, and worst fit cells. Each color corresponds to a label; solid lines are the fit, and dashed lines are the actual dynamics. Scatter plot summary for three cells with true *vs*. fitted observable reveals excellent agreement. (Right) Waterfall pot for the assessment of the convergence characteristics of the multi-start optimization for three cells.

To fit the model to single-cell data, we employed a multi-start local optimization approach. Specifically, we performed 1000 local optimizations, with initial points sampled randomly within parameter bounds.

The analysis of the computed parameter values revealed variability in the final objective function values across different cells, with the objective function value serving as an indicator of the goodness of fit (Figure 4B, left). This can be attributed to differences in the parameter vectors and noise realizations. Nevertheless, for all cells, we observed a good agreement between the data and model fit – as illustrated for a small subset of the observables (Figure 4B, center). This is especially noteworthy, as we still only have a single measurement per observable, but we were still able to estimate the time-resolved dynamics. Notably, the local optimization exhibited robust convergence, as evidenced by the prominent plateau in the waterfall plot (Figure 4B, right). Over 80% of the local optimization runs converged to the presumed global optimum for most cells. This suggested that the objective function landscape possesses only a tiny number of local minima with a substantial region of attraction and indicated that the number of optimization runs might be reduced if computational constraints arise.

In summary, the proposed parameter optimization pipeline yielded reliable fits for the rule-based model. This achievement is particularly noteworthy given that the model corresponds to an ordinary differential equation with 101 state variables and 40 estimated parameters, underscoring its robustness and potential applicability.

### Dynamic modeling allows for assessment of cell heterogeneity

Given that the proposed parameter optimization pipeline allows for reliable fitting of data, we assessed how well the parameters of individual cells and the parameter distribution on the population level can be reconstructed (Figure 5A). Here, we utilized the fact that the ground truth parameters are known for the considered synthetic data and examined the previously described scenario with three labels.

**Figure 5:**
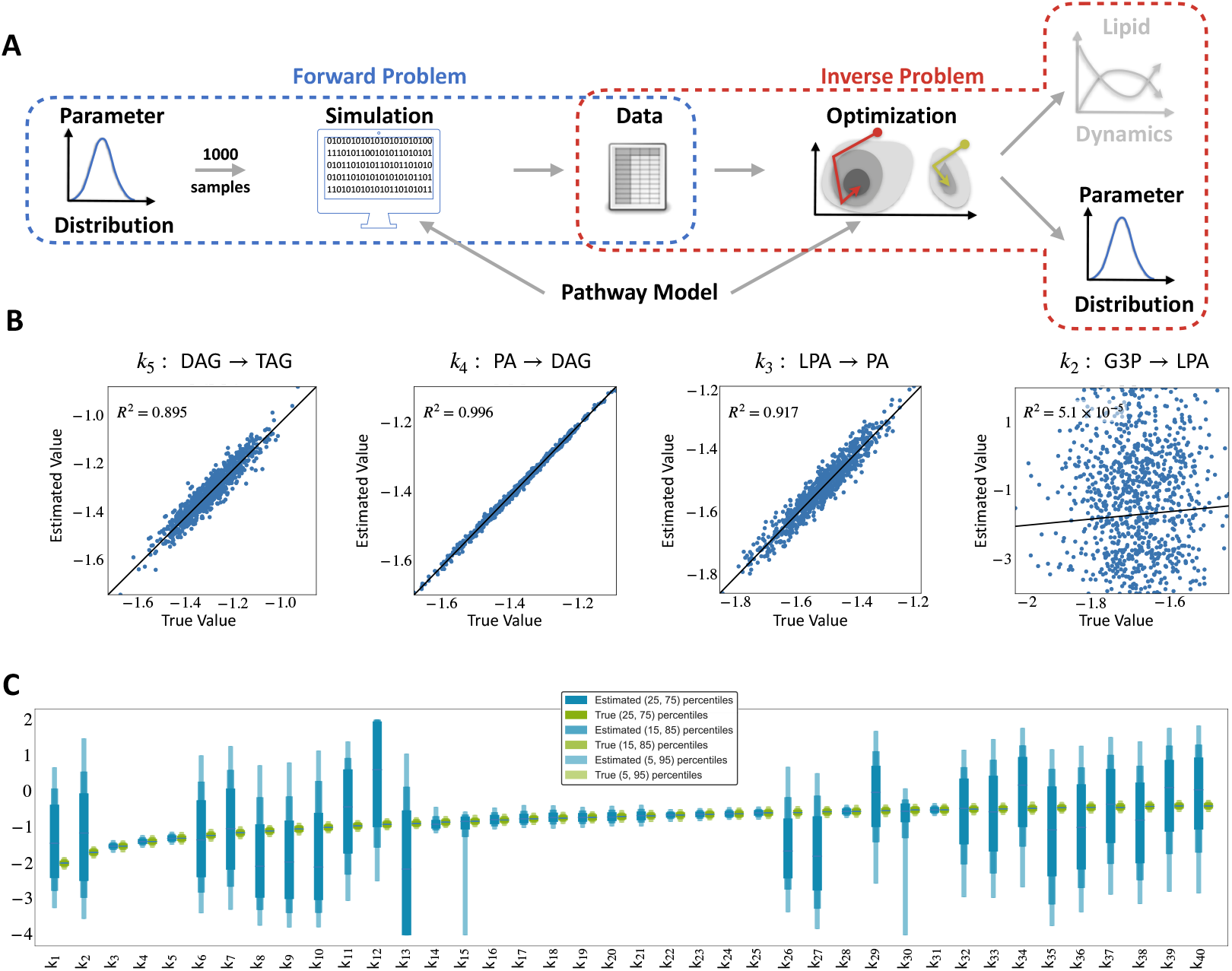
Analysis of single-cell parameters recovered from best local optimization. (A) Inverse problem with the focus on the recovery of parameter distribution. (B) Scatterplot for parameters of triglyceride synthesis (G3P → TAG). G3P is an unobserved state variable, while the other four metabolites are measured. (C) Boxplots of true and estimated parameter distributions for each parameter. The estimated ones are the best-found parameters for each cell after optimization. We see that the estimates of half the parameters have a good agreement (with some outliers on the 95 percentiles), while the others span multiple orders of magnitude.

To evaluate how well the parameters for individual cells were recovered, we compared the correlation between the maximum likelihood estimates and the ground truth values for each cell (Figure 5B). We found Pearson correlation coefficient ranging from −0.049 to 0.998. For 9 out of 40 parameters, a correlation above 0.8 was observed, while for 25 out of 40 parameters, the correlation was below 0.2. Interestingly, for the parameters with a low correlation, the estimated values spanned a much wider range, as indicated by the comparison of the empirical distributions of the maximum likelihood estimates and the ground truth distribution (Figure 5C). This indicates practical non-identifiability, which might be related to limited data or structural problems. Indeed, the uncertainties in the estimated parameters can be attributed to unobserved state variables, such as choline (C) and phosphocholine (C_PH_) (Supplementary Figure S3 A). These unobserved species are interconverted, and the parameters for the conversion rates (*k*_6_, *k*_32_) are not well determined (Figure 5C). Notably, the inspection of the multi-start optimization results confirms that there are multiple different optimization endpoints with similar objective function values. Thus, these endpoints appear to be part of a high-dimensional shape (Supplementary Figure S3 B-D).

In conclusion, the proposed experimental protocol with three labels in combination with the model-based analysis pipeline allows for the reliable assessment of a substantial fraction of the single-cell parameters. For those identifiable parameters, we were able to approximate the distribution across cells with a slight overestimation of the variance within the distribution.

### Quantifying the Impact of the Number of Labels on the Assessment of Cellular Dynamics

As the number of data points influences parameter identifiability and the quality of parameter estimates, we assessed the impact of the number of employed labels. We considered the use of zero to four labels, assuming that the unlabeled species are in steady state at *T*_0_ and that the labeling time points *T*_*i*_ are uniformly spaced between *T*_0_ and the measurement time *T*_*m*_ (Figure 6A). For each labeling setup, we generated synthetic data for 200 cells, using the same set of 200 sampled parameter vectors across all scenarios. We performed parameter estimation on each of the data sets as described above and quantified the impact of the number of labels on model complexity, computation time, parameter estimation, and parameter identifiability.

**Figure 6:**
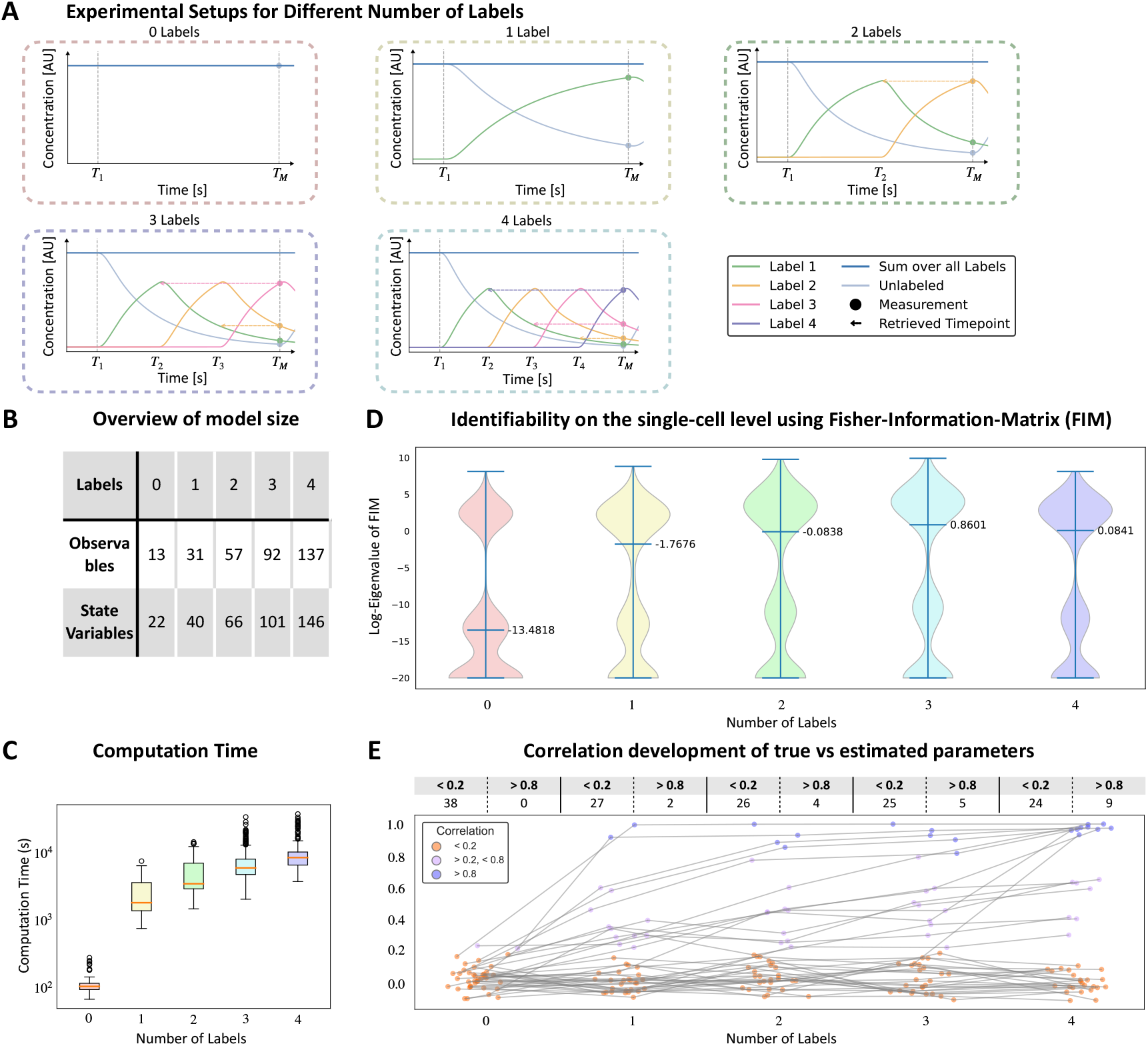
Impact of different numbers of labels on model size, computation time, and parameter identifiability. (A) Exemplary dynamics for each number of labels. The general experimental setup follows the same principle as Figure 2A. (B) Overview of the model size depending on the number of labels. (C) Average execution time of a complete multi-start local optimization for different numbers of labels. (D) Violin plots for eigenvalue spectra of Fisher information matrix. Each violin plot corresponds to a different number of labels used and plots the whole range of values, with the median highlighted separately. Eigenvalues smaller than 1 × 10^−20^ were set to 1 × 10^−20^ for ease of visualization. (E) Correlation of true vs. estimated parameters for each label. The table tracks the number of highly correlated (> 0.8) and uncorrelated (< 0.2) parameters.

The study of the model complexity for different labels revealed that the number of state variables and observables grows rapidly with the number of labels (Figure 6B), with a tenfold increase in observables from 0 to 4 labels. This increase is driven by the species with the highest number of label configurations, in this case, the TAG. The increased model complexity impacted the computation times required for optimization (Figure 6C), increasing by nearly two orders of magnitude (from 2 min to 170 min per local optimization). Importantly, for the rule-based implementation, which considers the equivalence of the labeling positions, the number of state variables and the computation time grow more slowly than the standard alternative.

To measure the influence of the number of labels on the identifiability of the parameter of individual cells, we used the Fisher information matrix. We computed the Fisher information matrix and its eigenvalue spectrum for the maximum likelihood estimates of the single-cell datasets for each number of labels. Subsequently, we compared the resulting spectra between the number of labels to assess how this number impacts identifiability. The eigenvalue spectrum of the Fisher information matrix provides information about the uncertainty of parameters (see, e.g., [Fröhlich et al., 2018a]) and hence about the information content of different experimental setups. The analysis of the eigenvalue spectrum showed that many eigenvalues are smaller than 10^−9^, indicating directions of uncertainty in the parameter space. The setup with zero labels had the most eigenvalues in this range and smaller eigenvalues overall (Figure 6 D). Incorporating additional labels shifted the eigenvalue distribution to larger values, signifying reduced uncertainty. The most significant shift occurred from 0 labels to 1 label. The increase from 1 label to 2 labels and from 2 labels to 3 labels resulted in an increase in the median by approximately one order of magnitude. Surprisingly, the increase from 3 labels to 4 labels did not add information, which is likely due to the equidistant spacing of time points. On a population scale, we investigated how increasing the number of labels affected the correlation between the estimated and true parameters (Figure 6 E). Here again, we saw a substantial shift between 0 and 1 labels. Additional labels resulted in a steady increase of highly correlated and a steady decrease in uncorrelated parameters, indicating higher overall accuracy of estimated parameters. In contrast to the individual scale, we observed continuous improvement of the correlation, even going from three to four labels.

The significance of the shift in both analyses from 0 to 1 label can be attributed to the qualitative transition from steady-state measurement to dynamic observation. As more labels are added, the impact becomes more quantitative rather than fundamentally changing the nature of the observed dynamics.

In summary, our investigation underscores the crucial role of labeling strategies in the study of cellular dynamics. The number of labels impacts computational demands as well as parameter estimation quality and uncertainty. An increase in label numbers decreased uncertainty at both individual and population levels. However, the benefit of additional labels tends to plateau after a certain point while the computational demands continue to rise. Therefore, determining the optimal number of labels requires balancing improved accuracy against available computational resources.

## Discussion

High-throughput single-cell experiments are primarily analyzed using statistical and bioinformatic approaches that provide limited information on cellular dynamics. In this study, we illustrated how cellular dynamics can be retrieved from high-throughput single-cell experiments when paired with our proposed experimental setup and analysis. We developed an integrated framework and outlined how the parameterization of cellular dynamics can be used to quantify the cell-to-cell variability of lipid metabolism.

Using synthetic data that resemble the key aspects of single-cell lipidomics measurements, we demonstrated that the proposed computational pipeline facilitates robust data fitting. In addition, a substantial fraction of the single-cell parameters was rendered identifiable. However, some parameters remained practically non-identifiable. As these parameters were mostly linked to unobserved states, this could be further investigated using structural identifiability analysis [Villaverde et al., 2018]. Complementary, experimental design could be used to optimize the information content about the identifiable parameters. A mere increase in the number of labels revealed limited information beyond a certain point – which might be due to the time-shifting characteristic – so an optimization of the timing and the use of non-equidistant labeling might be beneficial.

As we proved in the Supplementary that some measurements of different label combinations share a common dynamic, this theoretically allows us to use a much more simplified model that is label agnostic and incorporates the measurements as different time points instead of different label combinations. This would result in a drastically reduced model size but we would also lose many mixed labeled measurements. An investigation of trade-offs of these kinds of models could give further insights and guide the way for more efficient dynamic modeling.

The experimental setup, as we designed it, is intended for lipid metabolism. In principle, however, this approach can be generalized to other reaction networks if a set of labels can be found that satisfy the key assumptions of this setup. These key assumptions are as follows: the reactions must be agnostic to the labels, and the experiment must be able to start in a steady state. Without these conditions, tracing the dynamics of different labels back to a single one would be impossible. Additionally, we need an accommodating mathematical model that describes the process dynamics in the presence of multiple labels and accounts for the passing of a label from one metabolite to another. A prime example that could satisfy these conditions is the well-established 13C metabolic flux analysis (13C-MFA)[Ahn and Antoniewicz, 2011]. 13C-MFA, particularly nonstationary 13C-MFA (INST MFA), employs a similar concept by subdividing the dynamics. This allows for the measurement of dynamical changes within the labeled metabolites while the overall system remains in a steady state [Cheah and Young, 2018]. Applying our experimental setup to INST MFA is possible as it satisfies the necessary conditions. However, there are two issues: Firstly, while our study showed that even just one label could significantly improve the analysis, there currently is no other stable isotope for carbon aside from 13C. This could be tackled by using stable isotopes other than just carbon, but it would require a slightly different setup than described here since the dynamics would not be time-shifted versions of one another. Secondly, the positioning of the carbon atoms and other isotopes would need to be modeled in detail. This means that the overall number of labeling sites will, in general, be vast and thus significantly increase model complexity, a well-known problem for INST MFA [Wiechert and Nöh, 2013].

In conclusion, our new proposed experimental setup in combination with the appropriate mathematical modeling opens up new possibilities in regard to destructive measurement techniques, generating longitudinal data from a single measurement. As it is agnostic to the kind of label, it has the potential to be applied in a wide variety of fields in addition to single-cell lipidomics.

## Limitations of the study

The most limiting factor to this setup and why we resorted to synthetic data is the sensitivity of measurement devices. While the feasibility of single-cell lipidomics has been demonstrated [Thiele et al., 2019], it is not a standard approach yet and requires cells with high lipid content. It should be noted explicitly that our proposed experimental setup is currently theoretical in nature. As direct empirical measurements using the described single-cell lipidomics approach are not yet feasible due to limitations in measurement sensitivity, we resorted to synthetic data to demonstrate the potential of our framework. Despite this constraint, recent technological advancements [Zhang and Vertes, 2018, Li et al., 2021] suggest that such experiments will become practically viable in the foreseeable future. We, therefore, remain optimistic that the present theoretical setup will soon transition into experimental application, thereby providing deeper insights into cellular dynamics at single-cell resolution.

## Methods

### Mathematical Models

To model the lipid metabolism in a single cell, we use a system of ordinary differential equations (ODEs):

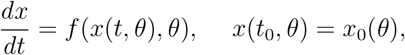

in which 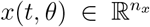 is the vector of state variables at time 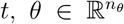 is the vector of parameters, and *f* the vector field describing the time evolution of the vector of state variables. The vector of state variables is at time *t*_0_ initialized with *x*_0_(*θ*). Throughout the study, we assume that this initial state is the equilibrium point for a single cell exposed to a label-free growth medium. Accordingly, the initial condition is defined as

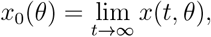

Metabolite levels are assumed to be partially observed (using, e.g., mass spectrometry). Metabolite levels are considered as sums over different lipid species (e.g., Triglycerides with different fatty acid chain lengths), only differentiating between the labeled fatty acids. The vector of observables at time *t* is denoted by 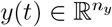,

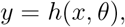

with observable mapping *h*. The observations are subject to measurement noise, which we consider here to be additive normally distributed. We denote all measurements with 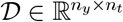,with

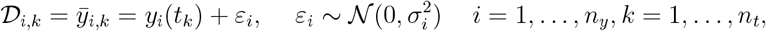

in which *n*_*y*_ is the number of observables, and *n*_*t*_ is the number of measurement time points.

In our model, state variables represent concentrations of metabolites. For each different combination of labels per lipid, one state variable is used. The considered molecules, as well as their respective number of state variables, can be found in TableS1. We consider mass action kinetics for each reaction and assume that reactions including labeled lipids have the same kinetic parameter for each label combination. The reactions and their corresponding rates are listed in TableS2.

### Synthetic Data Generation

To generate synthetic data, we sampled vectors of parameters from a pre-defined log-normal distribution. This distribution was chosen to ensure positivity in the parameter values. The means for each parameter individually were set to a value between 0.01 and 0.5. With a selected standard deviation of 0.1, we anticipated the samples of each parameter to encompass a range spanning between half and one order of magnitude. The sampled parameter vectors were used to simulate the mathematical model (Figure 4A, blue box).

For data generation, we simulated the ODE model using AMICI [Fröhlich et al., 2021], including a pre-equilibration step to mirror the experimental setup. To emulate real-world conditions, we introduced additive normally distributed measurement noise, with a standard deviation equal to 10 percent of the measurement value. The assumption of normality is commonly employed in systems biology studies Loos et al. [2018]. Indeed, the second most commonly used noise model is multiplicative log-normal noise, which, after transformation, yields on the log-transformed observable also additive normal noise Kreutz et al. [2007]. The generated data served as the starting point for parameter estimation (Figure 4A, red box).

For the initial analysis, we used the mathematical model with three labels. We generated a thousand parameter sets, with each set designed to represent an artificial cell. For each cell, we created a corresponding parameter estimation problem that we encoded in the PEtab format [Schmiester et al., 2021].

To investigate the effect of the number of labels on the parameter uncertainty (Figure 6), we used five different models, using between zero and four labels. Independent of the number of labels, the simulation time after pre-equilibration was kept constant. We generated 200 parameter sets in the same fashion as the previously generated ones. Within this set, we generated models and data for each label, resulting in five distinct datasets for each parameter set. This approach ensured consistency in parameter selection across different labels, allowing comparisons between the number of labels.

### Parameter estimation

To infer the unknown model parameters from data, we apply a maximum likelihood approach. Assuming independent, normally distributed measurement noise, the likelihood function is given by

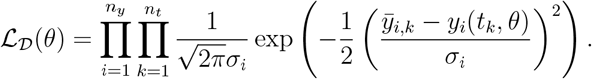

The maximum likelihood (ML) estimate is obtained by maximizing the likelihood function. For numerical stability and improved convergence, we minimize the negative log-likelihood:

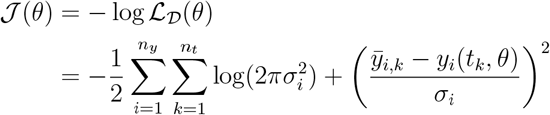

For the considered example, the parameters are constrained to the interval [10^−4^, 10^2^], spanning six orders of magnitude. We used multi-start gradient-based local optimization with 1000 log-uniformly distributed starting points for minimization. Optimization was performed in pyPESTO v0.3.1 [Schälte et al., 2023] using a PEtab [Schmiester et al., 2021] representation of our model and interfacing the Fides v0.7.8 [Fröhlich and Sorger, 2022] optimizer. The model was simulated using AMICI v0.18.0 [Fröhlich et al., 2021]. For a visualization of the workflow, we refer to Supplementary Figure S2.

### Uncertainty Analysis

To evaluate the impact of the number of labels on the identifiability of model parameters, we used the Fisher information matrix (FIM) at the maximum likelihood estimate. We computed the FIM via AMICI [Fröhlich et al., 2021] by summing over the product of the partial derivatives of the objective function with respect to the parameters, i.e., for any parameter 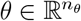

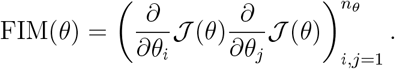

The eigenvalues of the FIM provide information about the likelihood function landscape that is close to the maximum likelihood estimate. Eigenvectors for large (positive) eigenvalues indicate weighted sums of parameters (directions in parameter space), which are well determined, while eigenvectors for eigenvalues that are zero indicate directions in which the likelihood function landscape is locally flat. Thus, the FIM at the maximum likelihood estimate provides information about the practical identifiability and sloppiness of the estimates. Accordingly, the eigenvalue spectrum can be used for uncertainty comparisons.

## Resource availability

### Lead contact

Requests for further information and resources should be directed to and will be fulfilled by the lead contact, Jan Hasenauer.

### Materials availability

This study did not generate new unique reagents.

### Data and code availability

The model, code for data generation, simulation, and inference, as well as the result files, are available on GitHub (
https://github.com/PaulJonasJost/Pseudo-time-trajectory-of-single-cell-lipidomics.git) and Zenodo (https://doi.org/10.5281/zenodo.12654308).

## Author contributions

Conceptualization, J.H. and C.T.; Methodology, P.J., D.W. and K.W.; Software, P.J.; Formal Analysis, P.J.; Writing – Original Draft, P.J., D.W. and J.H.; Writing -Review & Editing – all authors; Funding Acquisition, J.H.

## Acknowledgments

The icon in Figure 2 was created with BioRender.com.

