## Supplementary Information for "Pseudo-time trajectory of single-cell lipidomics: Suggestion for experimental setup and computational analysis"

### Proof of Time Shift

In this section, we will mathematically prove that under certain assumptions, a model that incorporates different labels at different time points allows us to obtain a pseudo-time trajectory.

#### Assumptions

We consider processes whose dynamics are governed by reactions with fluxes following the law of mass action kinetics. We note that Michaelis-Menten kinetics can be considered as well. While this case is not considered in this proof, we will outline how this can be derived afterward. As the reaction parameters are not the focus of this section, we shall slightly deviate from the notation in "Mathematical Models" in the methods section.

In general, we define a metabolic reaction network as a number of reactions $R_i$ between metabolites, i.e.

$$R_i: \sum_{j=1}^{|\mathcal{X}|} s_{i,j}^- X_j \longrightarrow \sum_{j=1}^{|\mathcal{X}|} s_{i,j}^+ X_j, \quad i = 1, \ldots, n_R, \tag{1}$$

in which $n_R \in \mathbb{N}$ is the number of reactions, $s_{i,j}^-$, $s_{i,j}^+ \in \mathbb{N}$ indicate the stoichiometry of reactants and products, respectively, $\mathcal{X}$ denotes the species considered in the reaction network and $|\mathcal{X}|$ the number of species considered. We only consider reactions up to the second order, as reactions of any higher order can either be written as chains of reactions up to the second order or can be neglected [Trautz, 1916]. Therefore, our reaction network is simplified to

$$R_i: aX + bY \longrightarrow \textit{products}, \quad a, b \in \{0, 1\},\ X, Y \in \mathcal{X},\ i = 1, \ldots, n_R, \tag{2}$$

Given a reaction network, its dynamics can be modeled with a system of ordinary differential equations with the following right-hand side,

$$\frac{dx}{dt} = f(x(t,\theta), \theta) = Sv(x(t,\theta), \theta), \quad x(t_0) = x_0, \tag{3}$$

in which $x(t,\theta) \in \mathbb{R}^n$ denotes the vector of state variables corresponding to different species, $S$ is the stoichiometric matrix, $v \in \mathbb{R}^{n_R}$ is the flux vector of the reactions and $\theta$ are the model parameters associated with the reaction fluxes. We shall denote a specific species with $X \in \mathcal{X}$ or $Y \in \mathcal{X}$ and write $\frac{dX}{dt}$ for its corresponding differential equation. Additionally, we assume that the system has a steady state, i.e.

$$\exists\, \bar{x}(\theta) \in \mathbb{R}^n \text{ s.t. } f(\bar{x}(\theta), t) = 0\ \forall t \geq 0$$

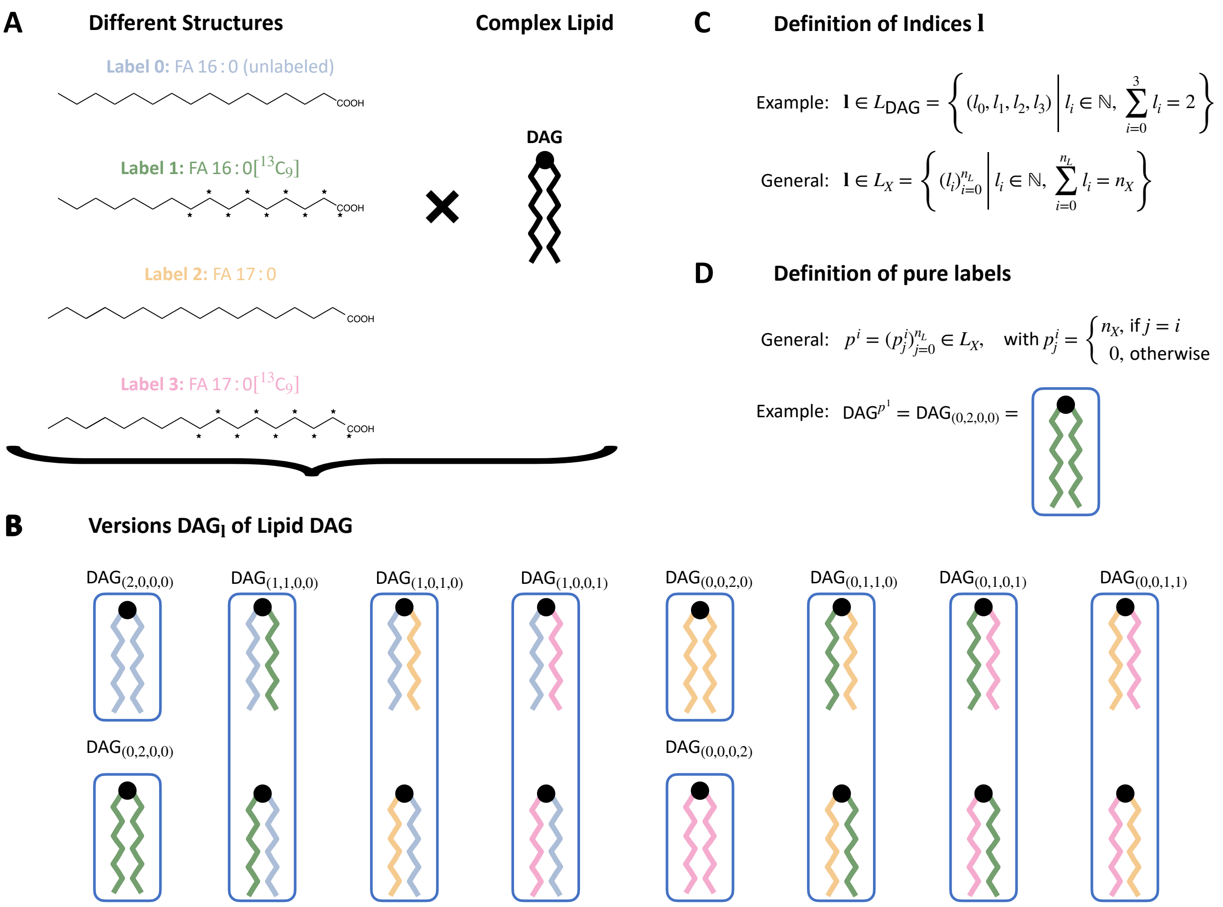


Figure S1: **Visual explanation of labeling and notation.** (A) We use different - but chemically similar - alkyne-fatty acids as metabolic labels. Therefore, any complex lipid (e.g., can now have multiple versions of itself depending on the label combination. (B) In the case of DAG, with three labels and one unlabeled, this results in 16 different versions. As we assume independence of the position, some versions are deemed equivalent in our model (blue boxes). Thus, the complexity of DAG is reduced to ten versions. (C) Definition of the index notation. Each entry $i$ in the vector index represents the number of labels $i$ in the specific variant of the DAG. They add up to the complex lipid's number of lipid binding sites. (D) For ease of notation, we also introduce the purely labeled lipids. $DAG^{p^1}$ represents the variant of DAG that has label 1 at each binding site.

We extend this model by incorporating a labeling strategy (Figure S1). With $n_L$, we denote the number of different labels $L_i$, $i = 0, \ldots, n_L$, in which $L_0$ represents the absence of a label. For each species $X \in \mathcal{X}$, we denote the number of labeling sites $X$ has with $n_X$. We define the species $X$ with a specific combination of labels $\mathbf{l}$ as $X^{\mathbf{l}}$, in which

$$\mathbf{l} \in L_X = \left\{ \mathbf{l} \middle| \mathbf{l} = (l_i)_{i=0}^{n_L} \in \mathbb{N}_0^{n_L+1}, \sum_{i=0}^{n_L} l_i = n_X \right\}, \tag{4}$$

is a multi-index from the set of possible multi indices $L_X$ for species $X$. Here, $l_i$ represents the number of labels $L_i$ the species $X^{\mathbf{l}}$ has. This notation has an implicit dependency on a species $X$, but as the multi-index is always used in combination with $X$, the context resolves this dependency.

Of specific interest are pure combinations that only consist of one label. For example, the species $X^{p^i}$ that only has labels of the kind $L_i$, i.e.

$$p^i = (p^i_j)_{j=0}^{n_L} \in L_X, \quad \text{with } p^i_j = \begin{cases} n_X, & \text{if } j = i \\ 0, & \text{otherwise} \end{cases} . \tag{5}$$

Given this setup, we assume the reactions to be independent of the labels:

**Assumption 1.** *Given a labeling defined in* (4) *and a reaction network, we combine them into a labeled reaction network in assuming the following*

1. *For each reaction* $R_i\,:\;\; aX + bY \longrightarrow$ *products, any labeling of* $X$ *or* $Y$ *can trigger reaction* $R_i$*, i.e. for all* $\mathbf{l} \in L_X$, $\mathbf{l}' \in L_Y$*, the reaction*

   $$R_i^{\mathbf{l},\mathbf{l}'}\,:\; aX^{\mathbf{l}} + bY^{\mathbf{l}'} \longrightarrow \textit{products}$$

   *is a valid reaction of the labeled metabolic reaction network.*

2. *The reaction rates are independent of the labels, i.e. given two reactions* $R_i^{\mathbf{l},\mathbf{l}'}, R_i^{\tilde{\mathbf{l}},\tilde{\mathbf{l}}'}$*, if the concentrations of* $X^{\mathbf{l}}$ *and* $X^{\tilde{\mathbf{l}}}$ *as well as* $Y^{\mathbf{l}'}$ *and* $Y^{\tilde{\mathbf{l}}'}$ *are equal, then the fluxes of both reactions are equal.*

In summary, Assumption 1.1 implies that a reaction can occur with any labeled version of its reactants. The labels on the products are dependent on the reactants. The second part assumes that reactions that only differ in the labeled versions of their reactants have the same reaction rates.

As per our experimental setup, the influx of the labels must be time-shifted versions of one another, except for label $L_0$.

**Assumption 2.** *Given a number* $n_L$ *of different labels* $L_i$*,* $i = 0, \ldots, n_L$ *that satisfy Assumption 1, we assume that for all reactions* $R_{L_i}$ *that represent an influx of label* $L_i$*,* $i = 0, \ldots, n_L$ *into the system, there is a corresponding reaction* $R_{L_j}$*,* $j = 0, \ldots, n_L$ *such that*

1. *The fluxes of those reactions can be described as time-shifts of one another, i.e.* $\nu_{R_{L_i}}(t) = \nu_{R_{L_j}}(t + T_{i,j})$*,* $T_{i,j} \in \mathbb{R}$*, for all* $1 \leq i, j \leq n_L$*, in which* $T_{i,j}$ *is the time-shift between labels* $L_i$ *and* $L_j$*.*

2. *The overall influx of all labels combined is constant, i.e.,* $\sum_{i=0}^{n_L} \nu_{R_{L_i}}(t) = C$*,* $C \in \mathbb{R}$.

Here, we assumed that the influxes of labels are time-shifted versions of one another and that the total influx is constant. Assumption 2.1 will ensure that the dynamics of the whole system are time-shifted of one another. Assumption 2.2 guarantees that the overall system remains in a steady state throughout the time course. An example would be the uptake of fatty acids in a cell, where we wash the cells at specific times and reapply a medium that only differs in the fatty acid content from the previous one.

#### Theorem and Proof

For the considered process, the following holds:

**Theorem 1.** *Given metabolic reaction network* (3) *with steady-state* $\bar{x} \in \mathbb{R}^n$ *and that Assumptions 1 and 2 hold, the following is ensured:*

1. *Given that we start in a steady state with only unlabeled metabolites, the sum over all labels of lipids is in a steady state, i.e.*

$$\forall X \in \mathcal{X}: \; \frac{d}{dt} \sum_{\mathbf{l} \in L_X} X^{\mathbf{l}} = 0$$

2. *For all* $X \in \mathcal{X}$*, the biochemical species that contain only one type of label are time-shifted versions of one another, i.e.*

$$\forall\, X \in \mathcal{X} \,\forall\, 1 \leq i, j \leq n_L : X^{p^i}(t) = X^{p^j}(t + T_{i,j}),$$

*in which* $T_{i,j} \in \mathbb{R}$ *is defined in Assumption 2.*

*Proof.* 1. We proof the first statement with a divide and conquer approach. We note that the ODE system stems from reactions, and thus, the differential equation for each species can be represented as a sum of reactions in which this species is involved. Additionally, we only consider reactions of up to the second order that follow mass action kinetics. Thus, it suffices to prove our point for each of those reactions individually. We will show that summing over the label combinations will yield the same ODE as the unlabeled reaction network for all three reactions. We use Assumption 1.1, by which reactions exist for all $\mathbf{l} \in L_X$ or none. We also make use of Assumption 2.2

(a) Zeroth order: By Assumption 2.2, the sum over any influx is constant. Thus, we can write straightforward

$$R_1 : \emptyset \longrightarrow X \;\Rightarrow\; \nu_1 = k_1$$

$$\sum_{\mathbf{l}\in L_X} \frac{dX^{\mathbf{l}}}{dt} = \frac{d}{dt}\sum_{\mathbf{l}\in L_X} X^{\mathbf{l}} = \sum_{\mathbf{l}\in L_X} k_1 = C \tag{6}$$

(b) First order:

$$R_2: \ X \longrightarrow Y \ \Rightarrow \ \nu_2 = k_2 X$$

We first take a look at the product site

$$\frac{d}{dt}\sum_{\mathbf{l}\in L_X} X^{\mathbf{l}} = \sum_{\mathbf{l}\in L_X} -k_2 X^{\mathbf{l}} = -k_2 \frac{d}{dt}\sum_{\mathbf{l}\in L_X} X^{\mathbf{l}} \tag{7}$$

Now we use the fact that $\sum\limits_{\mathbf{l}\in L_X} \frac{dX^{\mathbf{l}}}{dt} + \sum\limits_{\mathbf{l}\in L_Y} \frac{dY_{\mathbf{l}}}{dt} = 0$, and thus, we can follow the same for the product(s) $Y$.

(c) Second order:

$$R_3: \ X + Y \longrightarrow \text{products} \ \Rightarrow \ \nu_3 = k_3 XY$$

Without loss of generality, we look at $X$

$$\frac{d}{dt}\sum_{\mathbf{l}\in L_X} X^{\mathbf{l}} = \sum_{\mathbf{l}\in L_X} -k_3 X^{\mathbf{l}} \sum_{\mathbf{l}'\in L_Y} Y_{\mathbf{l}'} = -k_3 \sum_{\mathbf{l}\in L_X} X^{\mathbf{l}} \sum_{\mathbf{l}'\in L_Y} Y_{\mathbf{l}'} \tag{8}$$

Following the same argumentation as in the first order, we can deduce the same for $Y$ and the products.

Now we can combine those, and by defining $\mathbf{X} := \sum\limits_{\mathbf{l}\in L_X} X^{\mathbf{l}}$, we can see that the differential equations agree with the original unlabeled network. Thus, a steady state of the sum is also a steady state of the unlabeled metabolic network and will, therefore, stay in a steady state.

2. In the following, we prove the second part of the theorem by rewriting reactions and species by summing up all species with a specific number of labels $L_i$. This creates a system of equations for label $L_i$, $i > 0$. Since $i$ is arbitrary, we can conclude, in combination with Assumption 2.1, that these systems are time-shifted versions of one another. The purely labeled species are then only a special case.

   The amount of species $X$ that has exactly $m$ times label $i$ is defined by:

$$S(i, X, m)(t) = \sum_{\mathbf{l}\in L_X | l_i = m} m X^{\mathbf{l}}(t).$$

   We can retrieve $\frac{dS(i,X,1)}{dt}$ by rewriting reactions in terms of exchanging labels. Given a reaction $Y \longleftrightarrow X + Z$ with flux $kY$. Without loss of generality, let us assume $n_Y = 2, n_X = n_Z = 1$, in which $n_Y, n_X, n_Z$ are taken from (4). Let us now look at $S(i, Y, m)$ for differing values of $m$.

(a) $m = 0$, $m = 2$

$Y^{\mathbf{1}}$ without a label $L_i$ ($l_i = 0$) cannot pass on a label $L_i$ to $X$ and will always end up in $S(i, X, 0)$. Therefore, we end up with

$$\frac{dS(i,X,0)}{dt} = kS(i,Y,0) + R, \tag{9}$$

in which $R$ indicates the terms of $S(i, Y, 1)$ and $S(i, Y, 2)$ that are considered in the next cases.

(b) $m = 2$

$S(i, Y, 2)$ only has label $L_i$, therefore it will always pass on $L_i$ in an exchange. Thus

$$\frac{dS(i,X,1)}{dt} = kS(i,Y,2) + R. \tag{10}$$

(c) $m = 1$

Given that we have two labels in $Y^{\mathbf{1}}$ of which exactly one is label $i$, the probability of passing on the label to $X$ is 50%. Therefore, half of the fluxes from $S(i, Y, 1)$ go to $S(i, X, 1)$ and the other half to $S(i, X, 0)$. We thus can extend equations (9) and (10)

$$\begin{aligned} \frac{dS(i,X,0)}{dt} &= kS(i,Y,0) + 0.5kS(i,Y,1) \\ \frac{dS(i,X,1)}{dt} &= kS(i,Y,2) + 0.5kS(i,Y,1). \end{aligned}$$

Any other reaction will follow the same reasoning, slightly altered depending on $n_X$ and $n_Y$.

Since we can describe the system $(S(i, X, m))_{X\in\mathcal{X},m\le n_X}$ independently of other labels, the label $i$ is interchangeable. As all $S(i, X, m)$ with $m > 1$ are zero before the influx of label $i$, $\forall t < T_i, m \ge 1 : S(i, X, m) = 0$ and in those cases $S(i, X, 0)$ is equal for all $i$, and we additionally know from Assumption 2.1, that the influxes are time-shifted versions of one another, we conclude that

$$S(i,X,m)(t) = S(j,X,m)(t + T_{i,j}), \;\; 1 \le i,j \le N_L, \; X \in \mathcal{X}.$$

Now our second claim follows directly, since the purely labeled $X^{p^i}$ are directly proportional to $S(i, X, n_X)$, and thus,

$$X^{p^i}(t) = \frac{S(i,X,n_X)(t)}{n_X} = \frac{S(j,X,n_X)(t+T_{i,j})}{n_X} = X^{p^j}(t)(t+T_{i,j}).$$

$\square$

This proof is theoretically applicable not only to metabolic networks but also to other systems, as long as they satisfy the assumptions made.

Assumption 1 can be made, as in our case, the labels are chemically similar alkyne-fatty acids themselves, and therefore, we can make the reasonable assumption that they do not influence what kind of reaction can or cannot occur [Raclot, 2003]. Furthermore, we choose fatty acids of very similar chain lengths to ensure that the metabolism of those fatty acids is comparable. Regarding Assumption 2, we deem it justifiable, as we apply media with identical composition with the sole exception of the fatty acid being a different one. Therefore, the two points in Assumption 2 are satisfied as long as the uptake of fatty acids is equal for the differing labels, which we already assumed with Assumption 1. For the synthetic data generation, we consider the case of a highly dynamic range of the concentrations and, thus, a relatively large percentage of labeled vs. unlabeled metabolites. In less dynamic pools, these percentages can be lower, but the proof and the computational method presented here are independent of this.

### Supplementary Tables and Figures

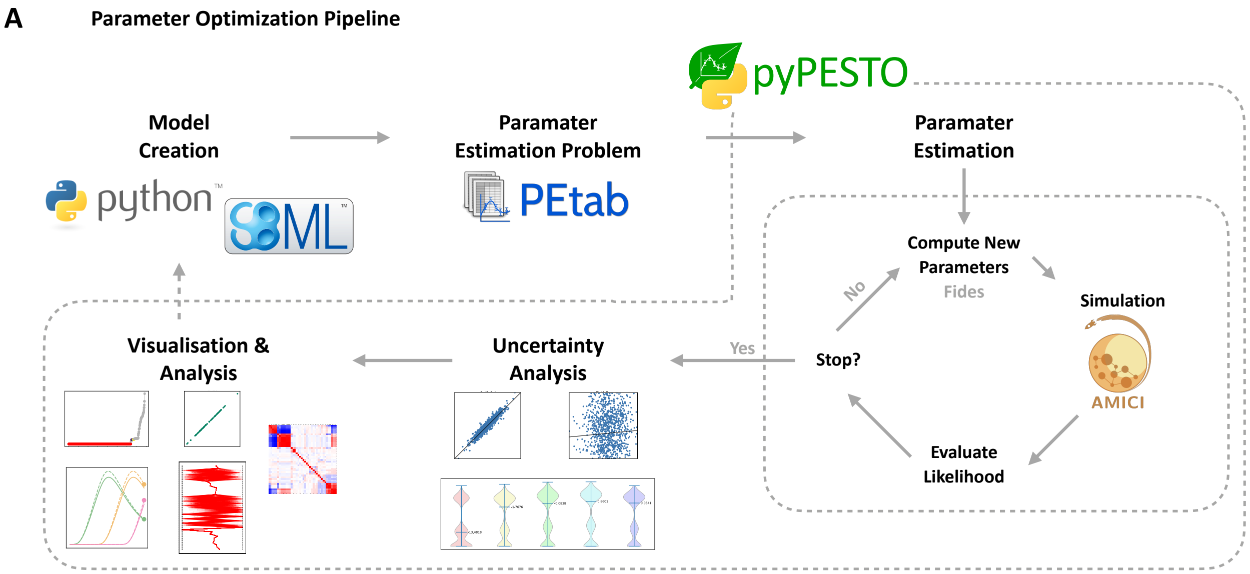


Figure S2: **Parameter Estimation Pipeline.** (A) The pipeline starts by creating an SBML model using a self-implemented rule-based model written in Python. Using the SBML model and the synthetically created parameters, PEtab [Schmiester et al., 2021] problems for each cell are generated. Subsequently, pyPESTO [Schälte et al., 2023] is used for parameter estimation, uncertainty analysis, and visualization. Parameter estimation is a cycle of simulation, likelihood evaluation, and new parameter calculation. For estimation, the Fides optimizer [Fröhlich and Sorger, 2022] is interfaced, and AMICI [Fröhlich et al., 2021] is used for simulation.

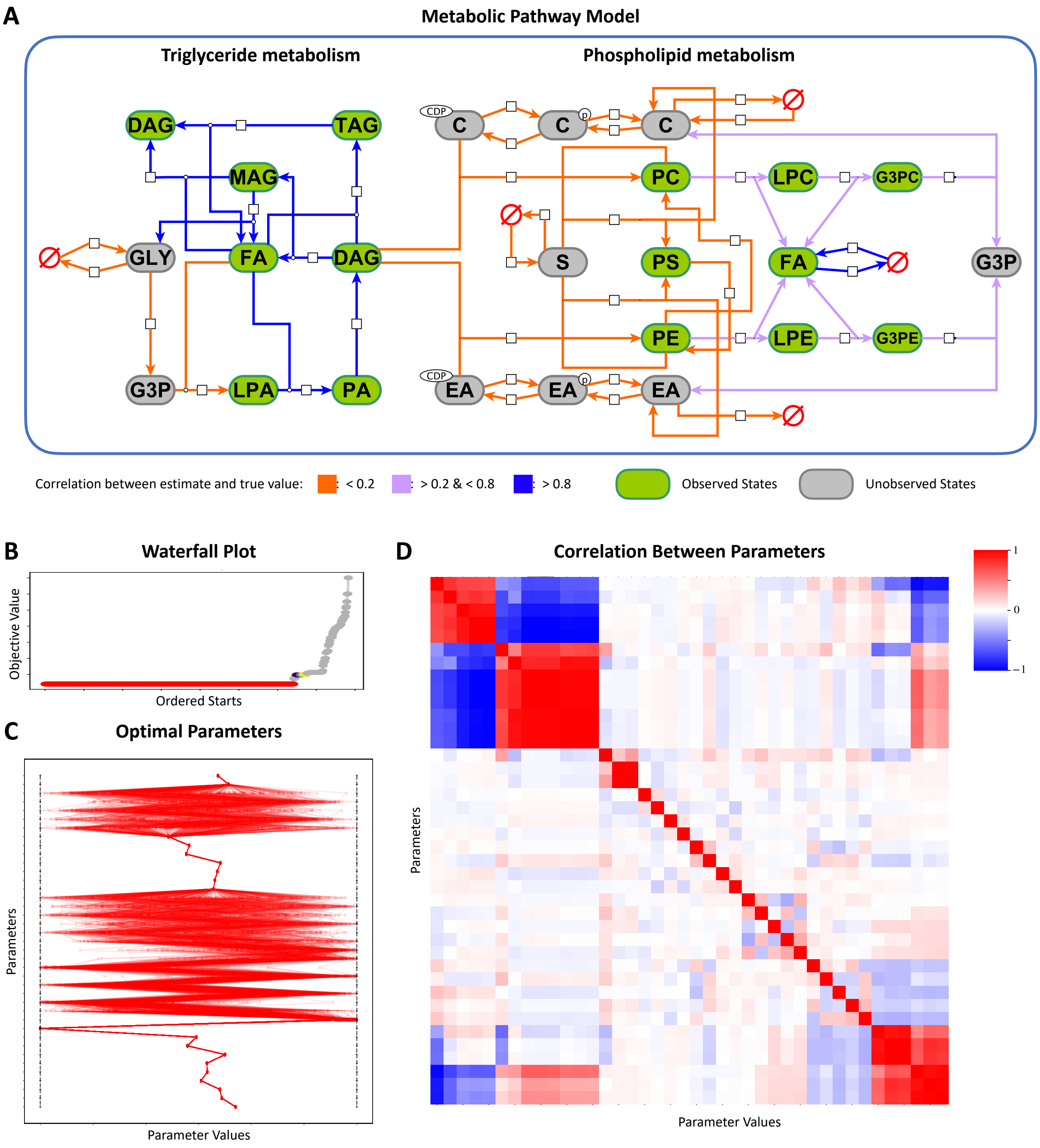


Figure S3: **Parameter uncertainties can be linked to unobservability.** (A) A graphical representation of the pathway model from Figure S1. Reactions are additionally colored by the correlation between the true and estimated parameter value of the corresponding parameter. (B) Waterfall plot of one multi-start local optimization. The final objective values of each start are sorted. Starts with the same color (except grey) correspond to almost equal final values. (C) Final parameter values of the best 400 starts plotted between the parameter bounds. All 400 starts had almost equal objective function values. (D) Correlation between parameters across the best 400 starts.

Table S1: List of the state variables considered in the mathematical model and their corresponding abbreviations.

| Name | Description |
|---|---|
| FA | Fatty acids |
| GLY | Glycerol |
| G3P | Glycerol-3-phosphates |
| LPA | Lysophosphatidic acids |
| PA | Phosphatidic acids |
| MAG | Monoacylglycerols |
| DAG | Diacylglycerols |
| TAG | Triacylglycerols |
| PE | Phosphatidylethanolamines |
| PC | Phosphatidylcholines |
| PS | Phosphatidylserines |
| EA | Ethanolamine |
| $\mathrm{EA}_{\mathrm{CDP}}$ | cytidine diphosphate (CDP) -ethanolamine |
| $\mathrm{EA}_{\mathrm{PH}}$ | Phosphoethanolamine |
| C | Choline |
| $C_{CDP}$ | CDP-Choline |
| $C_{PH}$ | Phosphocholine |
| S | Serine |
| LPE | Lysophosphatidylethanolamines |
| LPC | Lysophosphatidylcholines |
| G3PE | Glycero-phosphoethanolamines |
| G3PC | Glycero-phosphocholines |

Table S2: Reactions of the mathematical model. The first column indicates the rules used, i.e., the reactions of the base model without labels. In the second column, the corresponding rate laws - using the reactants in their base form - are listed. The third column shows how the number of reactions can be calculated through the metabolites participating in the reaction and their corresponding amount of labeling sites. The last column shows an example of the case of three labels. $n_L$ is the number of distinct labels used.

| **Rule** | **Rate Law** | **Number of Reactions** | |
|---|---|---|---|
| | | Formula | Example $n_L = 3$ |
| $\emptyset \longleftrightarrow FA$ | $k_{28} - k_{31}[FA]$ | $n_L + 1$ | 4 |
| $\emptyset \longleftrightarrow GLY$ | $k_{27} - k_{32}[GLY]$ | 1 | 1 |
| $GLY \longleftrightarrow G3P$ | $k_1[GLY] - k_{35}[G3P]$ | 1 | 1 |
| $G3P + FA \longrightarrow LPA$ | $k_2[G3P][FA]$ | $n_L + 1$ | 4 |
| $LPA + FA \longrightarrow PA$ | $k_3[LPA][FA]$ | $(n_L + 1)^2$ | 16 |
| $PA \longrightarrow DAG$ | $k_4[PA]$ | $\binom{2+n_L}{n_L}$ | 10 |
| $DAG + FA \longleftrightarrow TAG$ | $k_5[DAG][FA] - k_{24}[TAG]$ | $\binom{2+n_L}{n_L}(n_L + 1)$ | 40 |
| $DAG \longleftrightarrow MAG + FA$ | $k_{22}[DAG] - k_{23}[MAG][FA]$ | $(n_L + 1)^2$ | 16 |
| $MAG \longrightarrow GLY + FA$ | $k_{25}[MAG]$ | $n_L + 1$ | 4 |
| $\emptyset \longleftrightarrow C$ | $k_{36} - k_{37}[C]$ | 1 | 1 |
| $C \longleftrightarrow C_{PH}$ | $k_6[C] - k_{33}[C_{PH}]$ | 1 | 1 |
| $C_{PH} \longleftrightarrow C_{CDP}$ | $k_7[C_{PH}] - k_{38}[C_{CDP}]$ | 1 | 1 |
| $EA \longrightarrow \emptyset$ | $k_{40}$ | 1 | 1 |
| $EA \longleftrightarrow EA_{PH}$ | $k_8[EA] - k_{34}[EA_{PH}]$ | 1 | 1 |
| $EA_{PH} \longleftrightarrow EA_{CDP}$ | $k_9[EA_{PH}] - k_{39}[EA_{CDP}]$ | 1 | 1 |
| $\emptyset \longleftrightarrow S$ | $k_{26} - k_{29}[S]$ | 1 | 1 |
| $DAG + EA \longrightarrow PE$ | $k_{10}[DAG][EA_{CDP}]$ | $\binom{2+n_L}{n_L}$ | 10 |
| $DAG + C \longrightarrow PC$ | $k_{11}[DAG][C_{CDP}]$ | $\binom{2+n_L}{n_L}$ | 10 |
| $PC + S \longrightarrow PS + C$ | $k_{12}[PC][S]$ | $\binom{2+n_L}{n_L}$ | 10 |
| $PE + S \longrightarrow PS + EA$ | $k_{13}[PE][S]$ | $\binom{2+n_L}{n_L}$ | 10 |
| $PS \longrightarrow PE$ | $k_{14}[PS]$ | $\binom{2+n_L}{n_L}$ | 10 |
| $PE \longrightarrow PC$ | $k_{15}[PE]$ | $\binom{2+n_L}{n_L}$ | 10 |
| $PC \longrightarrow \emptyset$ | $k_{30}[PC]$ | $(n_L + 1)^2$ | 16 |
| $PC \longrightarrow LPC + FA$ | $k_{16}[PC]$ | $(n_L + 1)^2$ | 16 |
| $LPC \longrightarrow G3PC + FA$ | $k_{17}[LPC]$ | $n_L + 1$ | 4 |
| $G3PC \longrightarrow G3P + C$ | $k_{18}[G3PC]$ | 1 | 1 |
| $PE \longrightarrow LPE + FA$ | $k_{19}[PE]$ | $(n_L + 1)^2$ | 16 |
| $LPE \longrightarrow G3PE + FA$ | $k_{20}[LPE]$ | $n_L + 1$ | 4 |
| $G3PE \longrightarrow G3P + EA$ | $k_{21}[G3PE]$ | 1 | 1 |
